# Structural stability of Human serum albumin is modified in rheumatoid arthritis

**DOI:** 10.1101/2022.06.23.497357

**Authors:** Hsien-Jung L. Lin, David H. Parkinson, J. Connor Holman, W. Chad Thompson, Christian N. K. Anderson, Marcus Hadfield, Stephen Ames, Nathan R. Zuniga Pina, Jared N. Bowden, Colette Quinn, Lee D. Hansen, John C. Price

**Affiliations:** Department of Chemistry and Biochemistry, Brigham Young University, Provo, UT 84602 USA; TA Instruments, 860 W 410 N, Lindon, UT 84042 USA

**Author notes:** Authors contributed equally (Hsien-Jung L. Lin, David H. Parkinson). Corresponding author: John C. Price, 701E University Parkway, Provo Utah 84601.

**Keywords:** rheumatoid arthritis, serum albumin, differential scanning calorimetry, structure stability, proteomic, post-translational modification, protein surface accessibility

## Abstract

Differential scanning calorimetry (DSC) can interrogate changes in structure and/or concentration of the most abundant proteins in a biological sample via heat denaturation curves (HDCs). In blood serum for example, HDC changes are a result of either concentration or altered thermal stabilities for 7-10 proteins and has previously been shown capable of differentiating between sick and healthy human subjects. Here, we compare HDCs and proteomic profiles of 50 patients experiencing joint-inflammatory symptoms, 27 of which were clinically diagnosed with rheumatoid arthritis (RA). The HDC of all 50 subjects appeared significantly different from expected healthy curves, but comparison of additional differences between the RA the non-RA subjects allowed more specific understanding of RA samples. We used mass spectrometry (MS) to investigate the reasons behind the additional HDC changes in RA patients. The HDC differences do not appear to be directly related to differences in the concentrations of abundant serum proteins. Rather, the differences can be attributed to modified thermal stability of the most abundant protein, human serum albumin (HSA). By quantifying differences in the frequency of artificially induced post translational modifications (PTMs), we found that HSA in RA subjects had a much lower surface accessibility, indicating potential ligand or protein binding partners in certain regions that could explain the shift in HSA melting temperature in the RA HDCs. Several low abundance proteins were found to have significant changes in concentration in RA subjects and could be involved in or related to binding of HSA. Certain amino acid sites clusters were found to be less accessible in RA subjects, suggesting changes in HSA structure that may be related to changes in protein-protein interactions. These results all support a change in behavior of HSA which may give insight into mechanisms of RA pathology.

## INTRODUCTION

Rheumatoid arthritis (RA) is a systemic inflammatory autoimmune disease characterized by non-articular changes, symmetrical polyarthritis, and congenital symptoms (1, 2). Despite the prevalence of RA, the classification for the disease is considered definite only after the confirmed presence of chronic inflammation of the connective tissue in one joint, no reasonable alternative diagnosis, and scoring 6 or greater across the four different characterization domains (number of joints involved, abnormal antibody count, elevated acute-phase response, and duration of symptoms (3)). The late and tentative diagnosis of RA is mainly due to the elaborate, poorly understood etiology of the disease and a complex interplay between genetic and environmental factors. Effective RA management is correlated with early and aggressive treatment (2). Therefore, it remains crucial to develop an accurate, quick, and inexpensive way to diagnose RA, preferably without a tissue biopsy. The prognosis of RA patients depends heavily on early diagnosis since current treatments only relieve symptoms and slow progress, but do not cure the disease. Thus, the earlier the diagnosis, the better the prognosis for the patient.

Several low abundance proteins in human serum, such as C-reactive protein (4), rheumatoid factor (RF) (5), anti-citrullinated peptide antibodies (ACPA) (6), and anti-keratin antibody (AKA) (7) have been investigated for detection of pre-RA symptoms (5), but none have been found to serve as a biomarker for RA initiation. These proteins correlate with autoimmunity, but collectively make up an extremely minor percentage of human serum (<<1%), which often results in low sensitivity (8). Other metabolites, such as glucose (9), high-density lipoprotein (HDL) cholesterol (10) and vitamin D (11), have also been implicated in RA pathogenesis, but have not been used for diagnosis.

Current diagnostic tests for RA consist of measuring serum concentrations of rheumatoid factor (RF) and cyclic citrullinated peptide (CCP). Although RF concentration is widely used and the most accepted test for RA serologic diagnosis, it is not specific for RA (12). Elevated RF can be found in many other diseases, including other autoimmune diseases (Sjogren’s syndrome, systemic lupus erythematosus) (13, 14), chronic infections, cardiovascular disease, cancer, and normal aging (15). The sensitivity and specificity of RF for RA diagnosis are 62% and 89% respectively, and CCP’s sensitivity and specificity for RA diagnosis are 53-58% and 95-96% respectively (8). The RF and CCP tests are useful, but diagnosis often cannot occur until the disease has progressed significantly. Thus, more information about the causes of RA is needed to detect and intervene in RA development earlier and more accurately.

In this study, we compare RA-positive (RA) patients to RA-negative (non-RA) patients, all within a group of 50 who all came in for clinical testing because they were experiencing RA-like symptoms **(Figure 1)**. Comparing RA and non-RA subjects within a closed symptomatic group allows us to determine which proteomic changes come from RA-specific pathology, rather than generic inflammatory factors. We hope to reduce the confounding effect of comorbidities so that we can detect RA-specific differences. Here, we show our findings outlining RA-specific serum proteome changes and we propose a model for an RA-induced increase in HSA stability through potential binding partners, which we hope will offer valuable guidance in future research looking for better RA treatments and diagnostic models.

**Figure 1:**
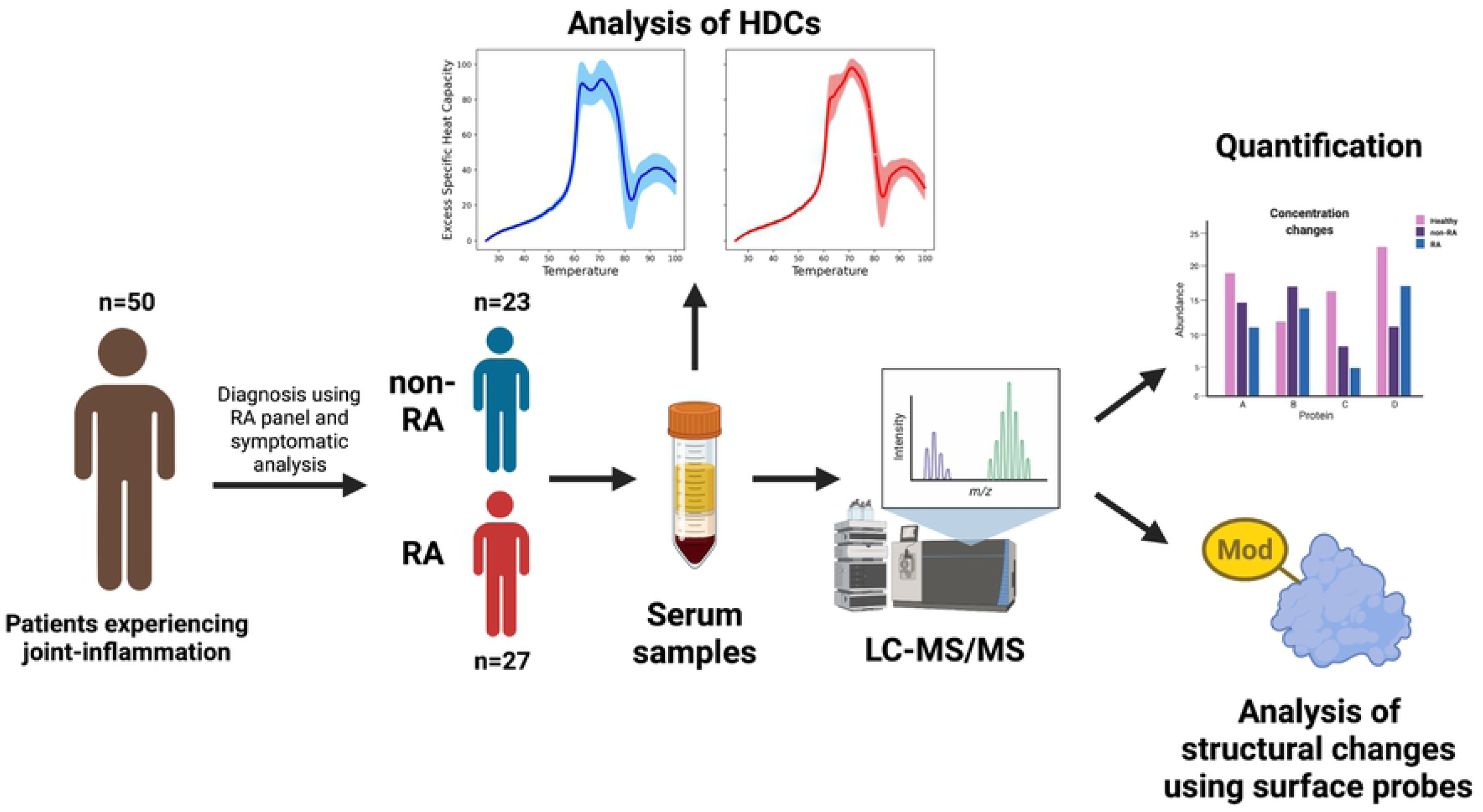
Experimental Flow. Blood serum samples from subjects that had physician ordered RA panels were used (n=50). Based on the clinical diagnosis, the samples are separated into an RA group (n=27) and a non-RA group (n=23). DSC was used to obtain the HDC, and the curve shift was compared between groups. LC/MS-MS experiments were performed to determine the mechanism behind the curve shift. Quantification and surface amino acid reactivity analyses were performed to determine significant differences in serum proteins between RA and non-RA groups.

A potential method to understand disease-specific pathology is from calorimetric thermograms (herein referred to as heat denaturation curves, HDCs) of patient serum obtained by Differential Scanning Calorimetry (DSC) (16–21). HDCs of serum from patients with several different diseases have been shown to exhibit reproducible shifts in the pattern of protein heat denaturation that are unique for those diseases (22–31). Although no mechanistic information is obtained, these differences must arise from changes in the concentrations and/or structures of the most abundant proteins (∼8 proteins) in the serum (32). In this study, HDCs were used to characterize the altered serum proteins in RA patients (clinically diagnosed according to symptoms and various biomarker levels). A characteristic HDC shift was seen across samples, and there was a significant relationship between RA diagnosis and HDC appearance. Mass spectrometry (MS) was then used to identify proteomic differences in the RA vs. non-RA samples, and we also looked for correlations between the proteome and HDC appearance (independent of RA diagnosis). Together, these results allowed us to understand which changes in the RA proteome could be attributed to the observed HDC differences.

We tested two possible mechanisms to explain this HDC shift **(Figure 2)**. First, changes in concentration of abundant proteins, such as human serum albumin (HSA), would alter the intensity of high abundance protein peaks, changing overall HDC shape (22, 33) **(Figure 2A)**. In this study, we focus primarily on HSA because it is the most abundant protein in plasma.

**Figure 2:**
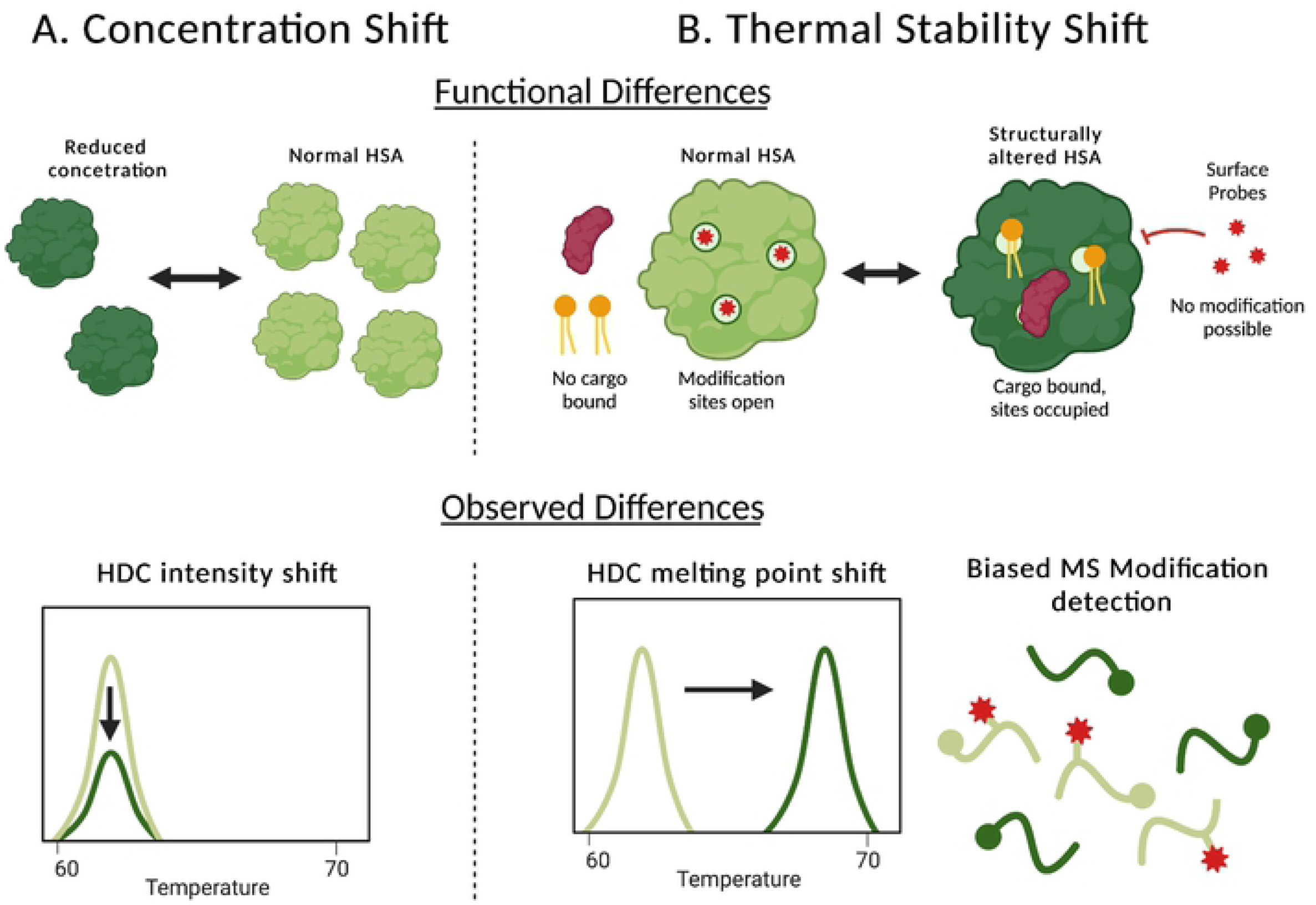
Two possible models explaining HDC shifts in RA subjects. The decreased peak ratio in RA samples could be explained by A) a relative HSA concentration decrease, reducing HDC signal intensity, or B) a shift in HSA thermal stability, shifting the HSA peak to the right, decreasing the first peak’s intensity, and increasing that of the second peak. This thermal stability could be the result of altered binding partners. The structural changes can be seen through biased detection of surface modifications on HSA. When cargo is unbound (top right, light green protein), binding sites are surface accessible for modification, when cargo is bound (top right, dark green protein), these sites are occupied, reducing surface accessibility and ability of these sites to be modified. These structural and functional differences at the molecular level could be the explanation for the observed shift in HDCs.

Second, HDC shape would be significantly altered by a change in thermal stability of abundant proteins, shifting their melting temperatures. For example, loading HSA with a fatty acid (octanoic acid) increases the melting temperature by 5 to 10 °C (34). Such changes in stability would most likely correlate with a change in these proteins’ tertiary structures and would be for a smaller portion of the total HSA (35) **(Figure 2B)**. We used MS to explore both concentration differences in HSA (and other abundant proteins), as well as HSA tertiary structure changes (by looking at surface reactivity (36, 37)) as potential causes of the characteristic shift in the HDCs. As outlined below, our data support a change in the HSA thermal stability **(Figure 2B)**. The MS-detected differences provide structural clues for HSA stability changes (38, 39).

## RESULTS AND DISCUSSION

We used a sample population of fifty anonymized serum samples from patients who experienced joint inflammatory symptoms. Patients ranged from age 12 to 88, with a median age of 50. Samples were not selected based on gender and the resulting sample set contained males (n=13) and females (n=37), matching statistical prevalence of RA (40). The rheumatoid arthritis panel (41) was conducted for the serum samples by ARUP Laboratories, including a rheumatoid factor (RF) and cyclic citrullinated peptide (CCP) test. Professional medical analysis of symptoms, paired with the CCP and RF levels, classified the 50 samples as coming from RA (n = 27) and non-RA (n = 23) subjects. **(Supplemental Data 1)**. Note that the serology results show only some of the factors used for RA clinical diagnosis. Other factors (joint involvement, acute phase reactants, and symptoms duration, etc. (42)) were used for diagnosis, but the supplementary diagnostic information was not provided for this study.

### Heat Denaturation Curves

Heat denaturation curves (HDC) were collected with a NanoDSC (TA Instruments, Lindon, UT). Forty-seven HDCs were obtained (HDCs for three subjects were uninterpretable due to errors during the sample injection). As seen in literature, HDCs for healthy subjects have two distinct peaks around 63 degrees and 71 degrees, which correspond to the known melting points for HSA and immunoglobulin proteins, respectively (30). The previous studies showed the low temperature peak at 63°C is primarily a combination of HSA and haptoglobin (HAPT), in which HSA dominates due to its much higher concentration (30). The high temperature peak at 71°C is primarily a combination of Immunoglobulin G (IgG) and Immunoglobulin A (IgA) underlain by the tail of the broad HSA peak (30). Healthy normal serum samples are reported in the literature with a more intense HSA (low temp) peak at 63°C and a comparatively lower intensity Ig (high temp) peak at 71°C (30). A separate study from Garbett et al. shows the 63 to 71 degree peak ratio in a cohort of healthy samples to be 1.59 ± 0.04 (22). As shown in **Figure 3A**, the non-RA HDCs show a decreased peak ratio (1.00 ± 0.23), and the RA HDCs show an even more substantial decrease in peak ratio (0.83 ± 0.16). A two-tailed t-test yields a p-value of 0.007 between these two groups, indicating that the HDC peak ratios are statistically different between RA and non-RA subjects **(Figure 3A, Supplemental Figures 1 and 2, Supplemental Data 2)**. This pattern is consistent with literature and can be seen in other auto-immune disorders such as lupus (23, 24). Similar to literature (25, 26), when the HDCs were ranked according to peak ratio (regardless of RA diagnosis), they could be separated into two groups that correlated with the RA diagnosis: low peak ratio (LPR, peak ratio < 1.00, n = 32), and high peak ratio (HPR, peak ratio > 1.00, n = 15) **(Figure 3B)**. Associating the HPR group with non-RA and the LPR group with RA gives a point-biserial correlation coefficient of 0.3966, meaning that 39.66% of the variability in peak ratio can be attributed to the RA diagnosis. With this association, a threshold ratio of 1.00 splits the samples (for the 47 HDCs obtained) with the smallest misclassification rate (27.7%). Using this threshold, 22 of the 32 samples (68.8%) in the LPR group are classified as RA while 12 of the 15 samples (80.0%) in the HPR groups are classified as non-RA. While 22 of the 27 RA samples (81.5%) are in the LPR group, and 12 of the 22 non-RA samples (54.5%) are in the HPR group **(Supplemental Data 1)**. These classification rates are likely impacted by the imperfect specificity and sensitivity of RA diagnosis mentioned earlier, as well as the presence of comorbidities in RA and non-RA subjects.

**Figure 3:**
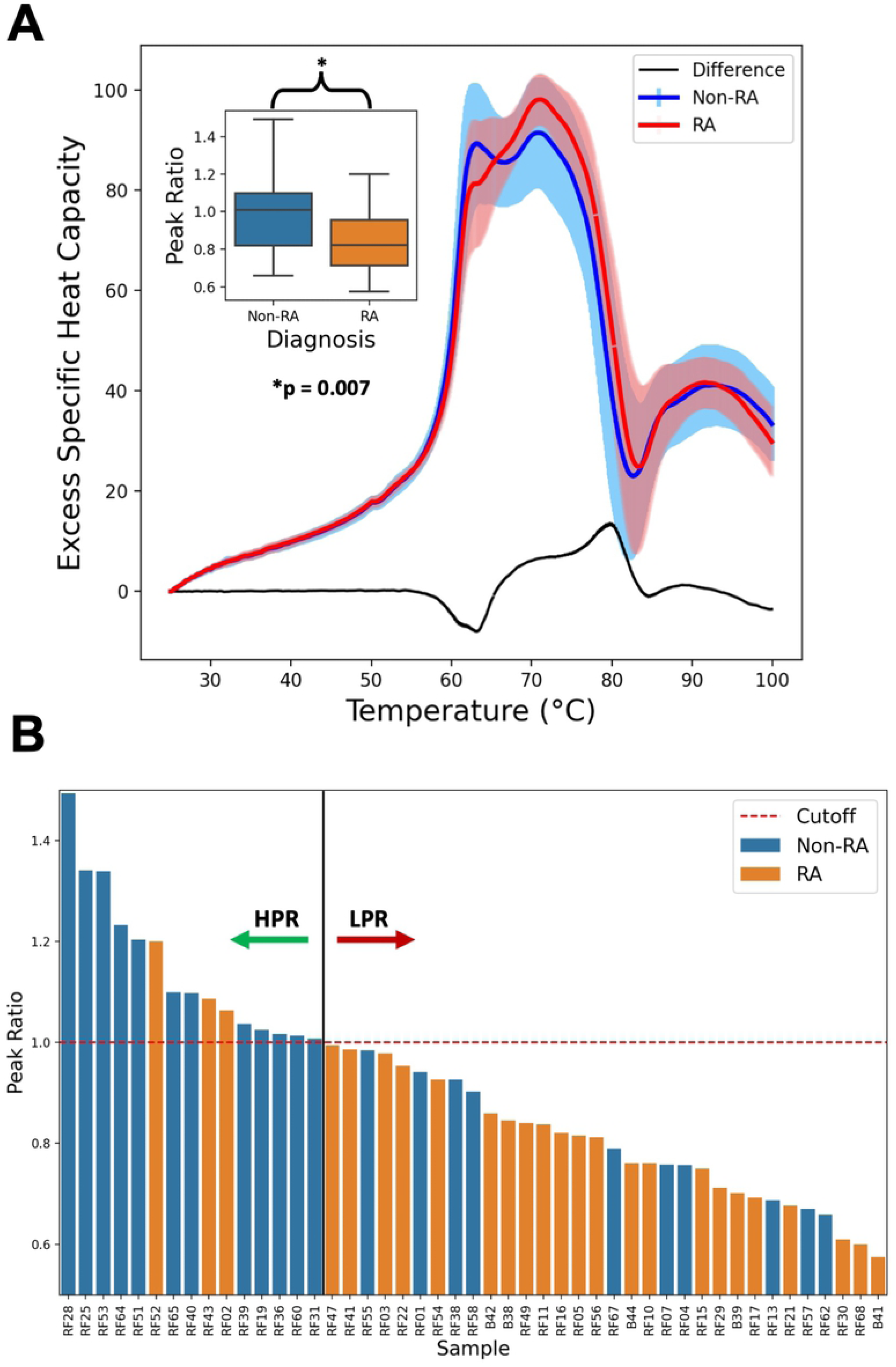
DSC results. This study focuses on the two peaks observed between 25 and 100°C of the heat denaturation curve. (A) The average HDC curve for non-RA and RA samples, with the difference between the two shown in black. The first peak from HSA is consistently found around 63°C (low temp peak) and the second Ig peak is always around 71°C (high temp peak). Inset for A shows the distribution of peak ratios from the HDC of RA and non-RA subjects. The difference in peak ratio between the non-RA and RA groups is statistically significant (p = 0.007) (B) The distribution of peak ratio of all samples, with a peak ratio threshold of 1.00 as the cut-off between the HPR and LPR groups.

Several of the non-RA samples are categorized in the LPR group, and this could be the result of other diseases or physiological differences (30) that alter HSA and other serum proteins, such as Lyme Disease, Lupus, or diabetes (22, 43). This seems likely given that all 50 subjects originally came in for testing because they were experiencing symptoms of discomfort and sickness. We are interested in mechanisms behind these HDC shifts **(Figure 2)**, so we used MS to evaluate the differences between both the RA/non-RA subjects and the HPR/LPR groups.

### Proteomics

#### Protein concentrations

The 50 serum samples were individually digested to tryptic peptides and analyzed using mass spectrometry to further explore the difference in protein content between RA and non-RA serum samples. Relative protein quantification analysis (PEAKS Studio_8.5, Bioinformatics Solutions Inc. (44), **Supplemental Data 3**) shows there are no significant differences in protein concentration between RA and non-RA groups or HPR and LPR groups for any of the top eight most abundant proteins (significant changes are defined as proteins with a fold change less than 0.5 or greater than 2 and a p-value less than 0.05) **(Figure 4, Supplemental Table 1)**. It is expected that specific autoantibody concentrations would increase in patients with RA (45–47), but since the RA antigen specific Ig population is a relatively small percentage of the entire Ig population, and significant sequence homology exists between immunoglobulins, it is difficult to distinguish target-specific antibodies using MS only. Also, the comparison was not against “healthy” controls, so that lack of significance in Ig could likely be because an Ig increase, non-specific to RA, may have occurred across many of the samples, elevating Ig levels altogether. These results suggest that a change in concentration of abundant serum proteins does not contribute to the decreased HDC peak ratio observed in RA samples.

**Figure 4:**
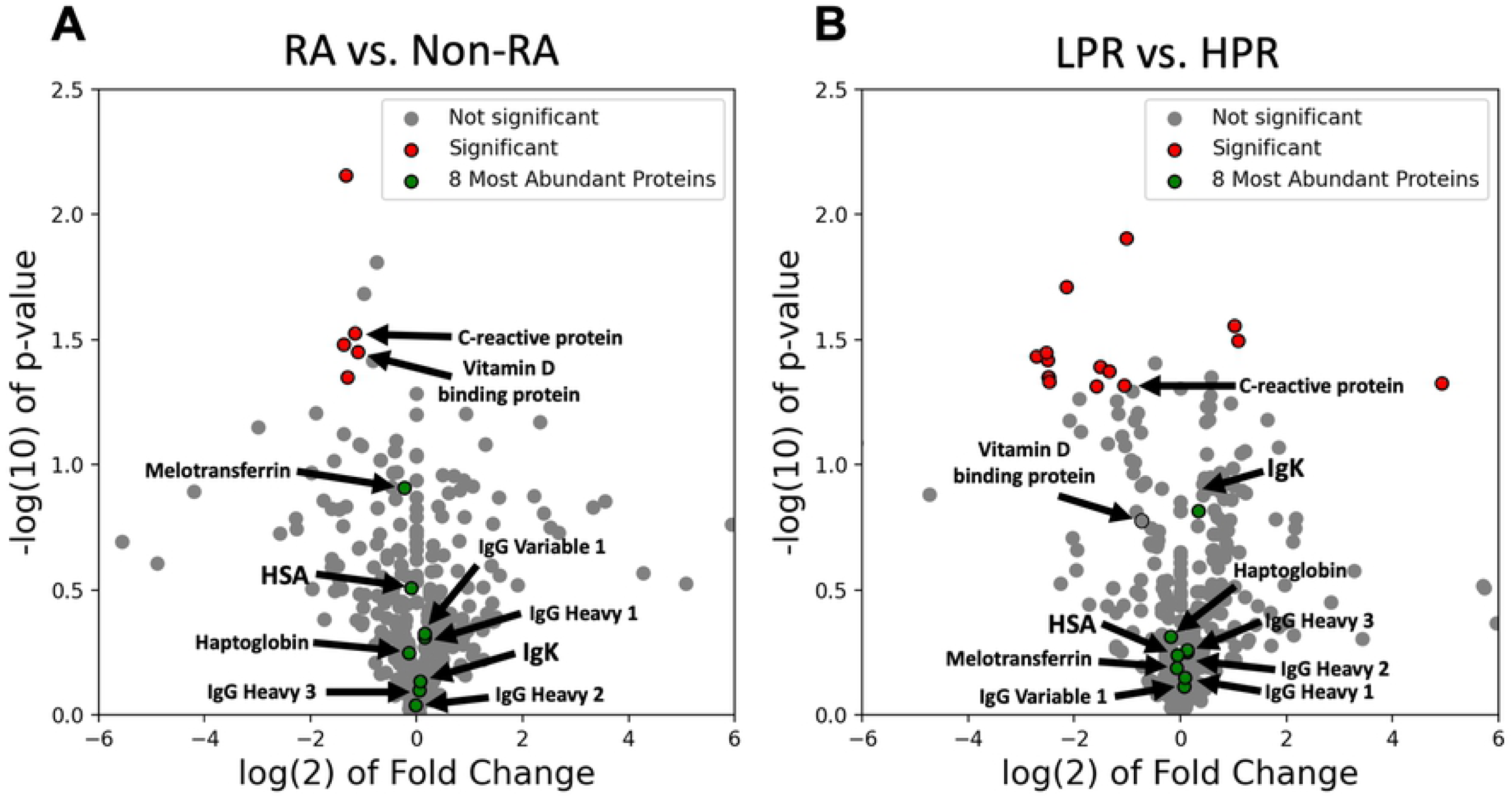
Protein Concentration differences in RA. Volcano plots indicating the fold change and p-value for all 421 detected proteins, comparing A) RA/Non-RA samples and (B) LPR/HPR samples. The top eight most abundant proteins are indicated in green (all insignificant), and the statistically significant proteins (−1 > fold change > 1, p-value < 0.05) are indicated in red. C-reactive protein is the only significant protein in both plots. The fold change is calculated in each comparison by dividing RA abundance by non-RA abundance and LPR abundance by HPR abundance.

Aside from the top eight proteins, overall proteomic analysis showed that among the 421 proteins compared, a statistically significant two-fold change was only seen in five proteins when comparing RA to non-RA samples **(Figure 4A)** and 14 proteins when comparing the HPR and LPR groups **(Figure 4B)** The only common significant protein between these two groups was C-reactive protein (CRP), a known RA biomarker. The significance of CRP in both the RA vs. Non-RA and HPR vs. LPR comparisons indicate that CRP may be involved in mechanisms accounting for the HDC shift seen in RA samples. It is important to note that CRP concentration is most likely upregulated for all subjects relative to healthy controls as has been described previously, but it is significantly lower in both LPR and RA groups. Vitamin D binding protein (VDBP, known to be related to RA (11)) was significantly downregulated in RA samples, and had no significant changes between HPR and LPR groups. These results suggests that although VDBP and other proteins may be associated with RA, their relatively low concentration means they are not directly affecting the HDC shift in RA samples. However, the change in the concentration of these proteins may affect our measurements because they have interactions with the very abundant proteins measured in the HDC **(Figure 2B).**

#### Structural changes in high abundance proteins

Since protein concentration doesn’t directly account for the difference in HDCs between RA and non-RA samples, and there is also no link between concentration and the HPR and LPR groups. Therefore, we expect, similar to other diseases explored in literature, that the observed HDC shifts among RA patients and the LPR group are caused by changes in thermal stability for one of the most abundant serum proteins. We simulated shifts in the melting temperatures of various percentages of each of the top eight serum proteins (using individually measured HDCs of these abundant proteins from literature (22)) could recapitulate the observed changes. We found an increase in the melting temperature for a small fraction (∼10%) of the HSA pool could explain the observed HDC shift **(Supplemental Figure 3)**. Changes in HSA melting temperature could result from new ligand binding, protein interactors, or tertiary structure (48–50). Here, we tested for structural changes of HSA through analysis of covalently modified amino acid profiles between the RA and non-RA samples. Both biological and artificially induced modifications were considered. Changes in biological modifications could show altered RA biochemistry, and changes in artificially induced modifications would show variations in surface accessibility of certain regions of a protein. If RA-specific protein conformation changes are responsible for changes in the HDCs, we also expect these amino acid modification (AAmod) profiles between RA and non-RA groups, to be correlated with the observed HDC groups (HPR and LPR).

Protein Prospector (UCSF) and PEAKS studio (Bioinformatics Solutions Inc) were both used for contrasting analysis of the PTM data. Multiple peptide modifications were observed as noncanonical m/z shifts with Protein Prospector, including a modification of +183 m/z, which was the most frequently observed modification (41 peptides) on HSA **(Supplemental Data 4)**. PEAKs Studio’s analysis of HSA proteins and each AAmod confirmed the +183 m/z modification as an aminoethylbenzenesulfonylflouride modification (AEBSF) which came from the protease inhibitor cocktail we added before processing the serum. Thus, AEBSF was an artificially induced, non-biological PTM. HSA had 185 modified sites that were observed in more than 12 of the samples. Of the 185 total AAmod sites on HSA, there were 33 observed modification types, with the top ten most frequent being AEBSF, 41; Dehydration, 28; Hexose, 17; Deamidation, 14; Iodination, 14; Oxidation, 9; Citrulline, 8; Formylation, 5; Amidation, 5; and Di-iodination, 4. 71% of these AAmod sites are specific for only one type of modification **(Supplemental Data 4)**. AEBSF was the only AAmod that showed statistically significant differences between RA and non-RA groups **(Supplemental Data 5)**. Since the AEBSF modification was synthetically introduced, it is not causing the change in HSA structure but is reporting the fact that the in vivo structure was changed for these reactive sites. Non-RA subjects have, on average, 1.9 times more AEBSF modifications that RA subjects (p = 0.023). Since there were significantly fewer AEBSF modifications in RA subjects, it suggests that AAmod sites are less accessible in RA HSA, suggesting conformational changes or a potential increase in binding partners in RA HSA.

#### AEBSF as a probe of surface reactivity

AEBSF is an irreversible serine protease inhibitor which can react with surface accessible nucleophilic amino acids such as Serine (S), Lysine (K), Tyrosine (Y), Histidine (H), and the amino-terminus **(Figure 5A)** (51, 52). Like other good surface modifiers (diethyl pyrocarbonate (37) or diazonium salt (36)), we can use its prevalence to identify changes in surface area accessibility of individual amino acids on proteins between samples. The AEBSF modifications observed on HSA were most frequently observed on lysine (28 different lysine residues), tyrosine (9 different residues), serine (2 different residues), and histidine (2 different residues) **(Supplemental Data 5)**.

**Figure 5:**
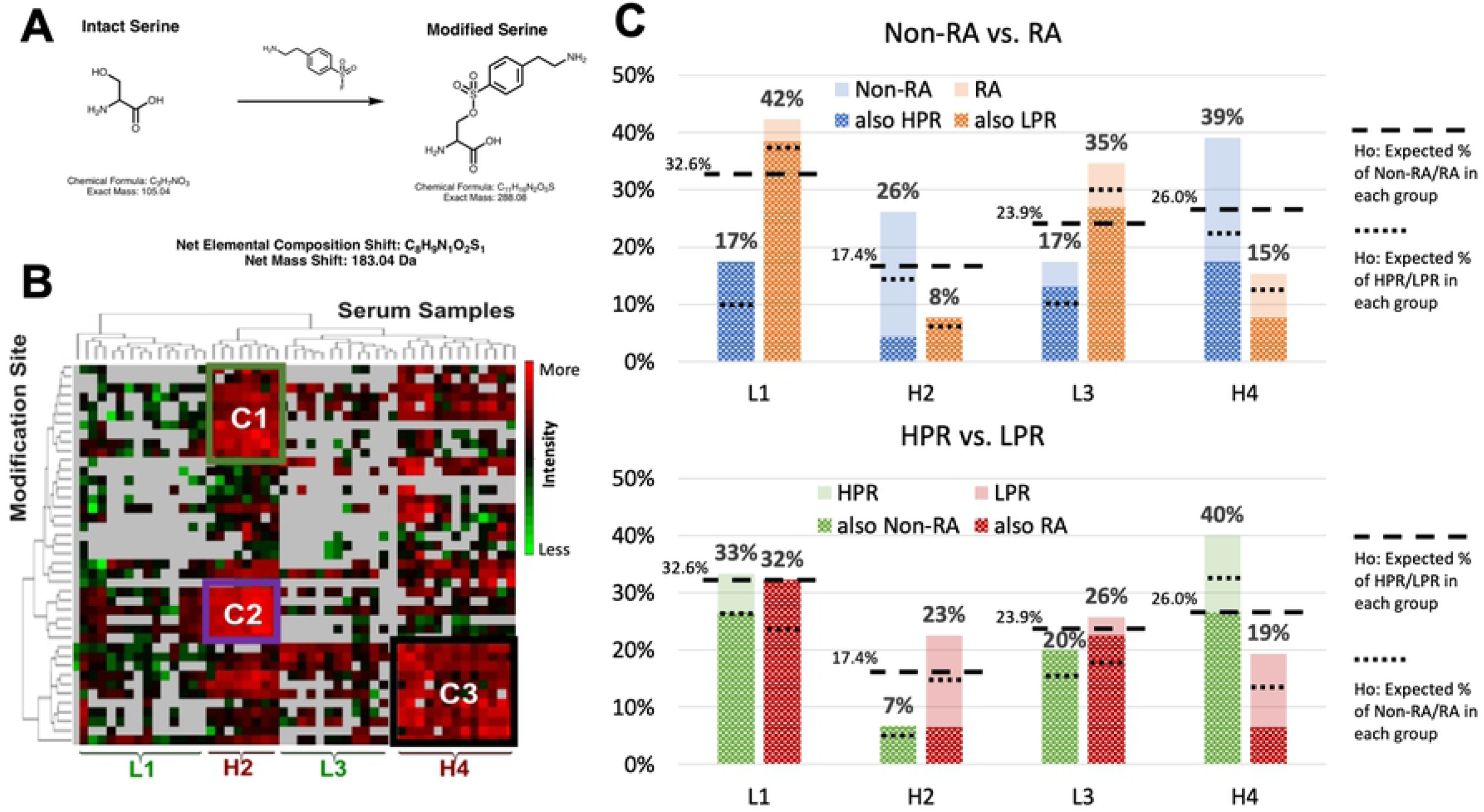
AEBSF modification of HSA. **(A)** The chemical reaction of the AEBSF modification on Serine. The reaction is similar for other nucleophilic amino acids. **(B)** The heatmap generated with PNNL Inferno showing the intensity differences of AEBSF modification at different HSA sites between different samples. The AEBSF modification amino acid number for HSA is listed on the y-axis, and the serum sample number on the x-axis. The samples are separated into 4 groups according to the hierarchy branch of serum samples, from left to right **(Supplemental data 5)**. Group L1 and L3’s AEBSF modifications are less intense than group H2 and H4 (L stands for lower intensity and H stands for higher intensity). The three clusters, C1 (green), C2 (purple), and C3 (black), are the most intense AEBSF modification clusters and are examined to characterize the modification further. **(C)** The bar graph shows the number of RA/non-RA and HPR/LPR samples expected in each AEBSF modification group (L1, H2, L3, H4). The percentage of RA samples in L1, H2, L3, H4, is 73%, 25%, 69%, and 31% respectively. For LPR, it is 67%, 89%, 73%, and 50%, respectively.

To visualize patterns in AEBSF modification between samples and across AAmod sites, we used PNNL Inferno (53) to generate a hierarchical grouped heatmap from the patient-specific ion intensities for each modification site **(Figure 5B)**. From our MS data, samples were sorted into clusters with serum samples on the horizontal axis and HSA modification sites on the vertical axis. The hierarchical order separated the samples into 4 groups. Of the 4 groups, two groups have higher signal intensity (H2 & H4), and two groups have lower signal intensity (L1 & L3). H2 has higher signal intensity at sites shown in clusters C1 and C2, H4 has higher signal intensity at the specific sites in cluster C3, and groups L1 and L2 have lower signal intensity across all sites. Higher signal intensity indicates a greater level of AEBSF modification. Each of these four clusters (L1, H2, L3, and H4) are made up of 32.6%, 17.4%, 23.9%, and 26.0% of the serum samples, respectively. Based on the number of samples, the clinical assessment and our HDC results can anticipate how random assignments would group the samples (null hypothesis, H_o_, **Figure 5C**). Therefore, we would expect 32.6%, 17.4%, 23.9%, and 26.0% of the samples in each of the RA/non-RA/HPR/LPR groups to be present in each of these four clusters. As shown in **Figure 5C**, we found that a much greater proportion of the RA samples were found in the L1 and L3 groups (42% and 35%, respectively) compared to the non-RA samples (17% and 17%, respectively). A lower percentage of RA samples were in the H2 and H4 groups (8% and 15%, respectively) compared to the non-RA samples (26% and 39%, respectively). The L1 and L3 groups contained close to the expected proportion of samples from the HPR and LPR groups, but in the H4 group (containing a high proportion of non-RA samples), we saw a higher-than-expected percentage of HPR samples (40% of HPR samples were in the H4 group, compared to 19% of LPR samples). However, in the H2 group (also containing a high proportion of non-RA samples), we saw a higher-than-expected percentage of LPR samples (23% of LPR samples compared to 7% of HPR samples). This suggests that the high AEBSF frequency at the AAmod sites in clusters C1 and C2 are connected to a decrease in HDC peak ratio but an RA-negative diagnosis. In fact, more intense AEBSF modifications at these C1 and C2 sites may give insight into why certain non-RA samples exhibited a low HDC peak ratio (increased surface accessibility from other factors not specific to RA). On the other hand, the high AEBSF frequency at the modification sites in clusters C3 are connected to both a higher HDC peak ratio and non-RA subjects **(Figure 5B,5C)**, indicating that decreased accessibility of the C3 amino acid binding sites seen in RA samples may be directly linked to the observed HDC shift seen in RA samples.

Together, this pattern suggesting that HSA in the RA/LPR groups may have binding partners or other ligand interactors that block those C3 sites. Additionally, the association between HPR and RA samples in the H2 group suggest that the decreased accessibility of C1/C2 AAmod sites, likely due to binding partners or other conformational changes, are unlikely to be the cause of the increased HDC shifts observed in RA HSA. These binding partners could be related to other diseases that the RA-negative (yet still discomforted) patients were experiencing when they came in to be tested for RA.

It should be noted that the clustering in our heatmap in **Figure 5B** is data-driven using these 50 subjects as a training set. Therefore, statistical inference, error bars, and p-values are not appropriate as we analyze how the data in **Figure 5C** deviates from our null hypothesis. To test the hypothesis that these patterns can be applied to a population with statistical confidence additional groups of non-RA and RA would need to be collected and compared to our clustered model. Even without this statistical assessment, previously published literature gives us valuable insight into these AEBSF groups and the modification sites in relation to specific changes in HSA tertiary structure.

#### Potential Binding Surfaces on HSA

The 3-dimensional structure of HSA has three recognized domains, with two subdomains each (54). There are also nine known binding pockets distributed throughout the three domains. Two drug binding sites, Sudlow sites I and II are located in domains IIA and IIIA (55, 56), respectively **(Figure 6B**). The HSA structure and the AAmod sites for each of the three clusters was visualized with UCSF Chimera (version 1.15) (57), with C1 sites in blue, C2 sites in red, and C3 sites in green **(Figure 6A)**. AEBSF modification sites in C1 and C3 are mostly in domain II: 70% and 55%, respectively. AEBSF modification sites in C2 and are mostly in domain I (50%) **(Figure 6C, Supplemental Data 5)**. Sudlow Site I (IIA) has the most frequently observed (33%) AEBSF modification sites from all three clusters combined **(Figure 6B, Supplemental Data 5)**. The modified amino acids in C1 and C3 are mostly lysine, and mostly tyrosine in C2 **(Figure 6D, Supplemental Data 5)**. PyRosetta (58) was used to extract the secondary structure and surface accessible surface area (SASA) scores from a representative crystal structure of HSA (PDB ID: 1N5U (59)); 81% of the modification sites are on an α-helix, and 19% are on a loop **(Figure 6E, Supplemental Data 5)**. The average SASA scores of C1, C2, and C3 are 95.9 ± 37.2, 37.1 ± 32.5, and 81.8 ± 37.3 **(Figure 6F, Supplemental Data 5)** where a larger value indicates more surface accessibility.

**Figure 6:**
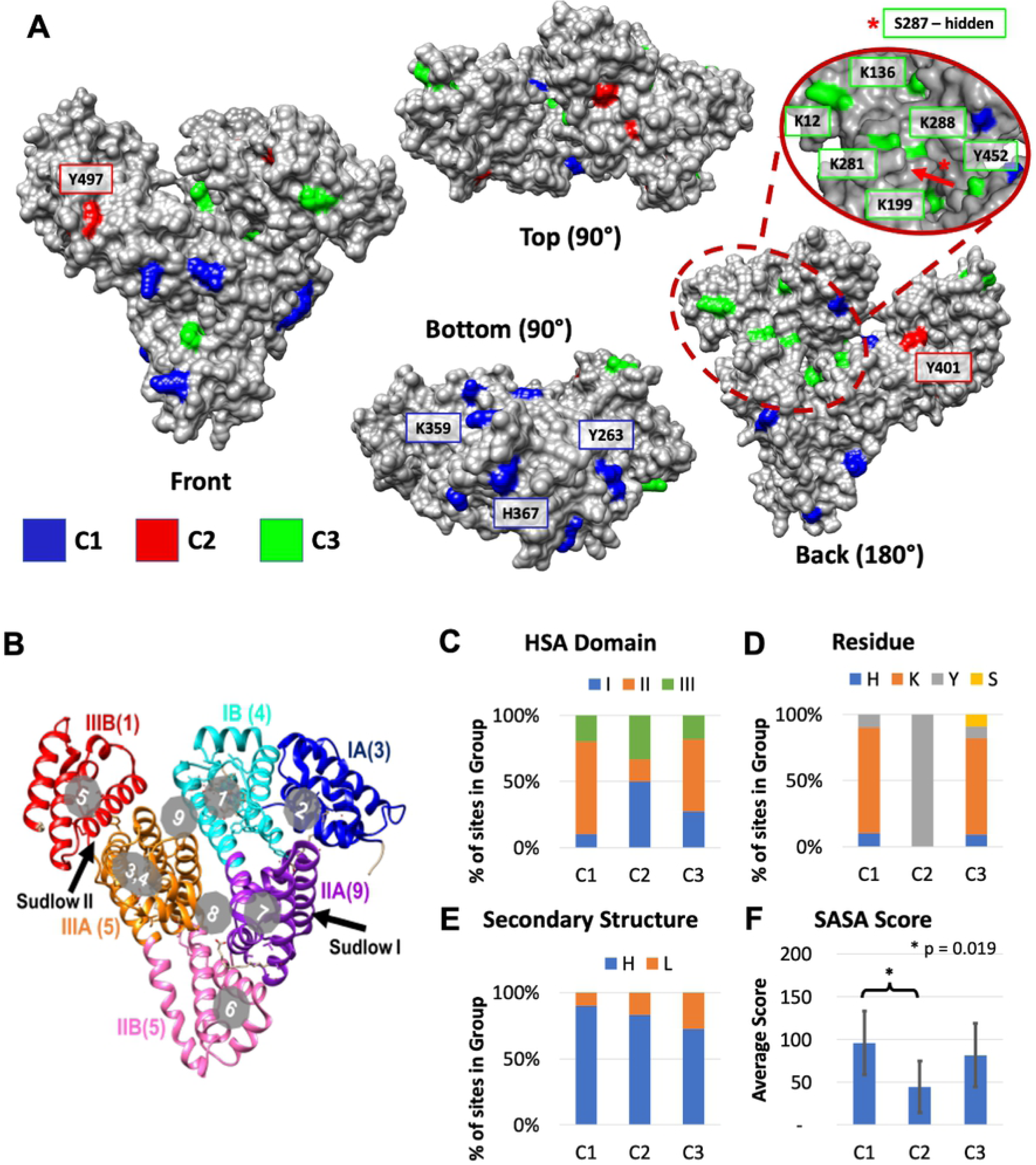
Characterizing AEBSF modification site of the 3 clusters (C1, C2, C3) on HSA. **(A)** A representative HSA crystal structure (PDB ID: 1N5U) with the 3 AEBSF modification sites clusters colored. The three clusters, C1, C2, and C3 are colored in blue, red, and green, respectively. Individual C1 and C2 sites that are significantly less accessible in RA HSA are labeled. The red oval indicates a C3-rich region in domain I that could be a plausible binding site for RA-specific interactors that most likely to increase HSA stability. **(B)** The HSA structure is colored by its 3 domains (I, II, III), and subdomains (A, B). The number of AEBSF modification sites regardless of cluster designation in each subdomain is listed in parentheses. The 9 known cargo binding pockets are shown in the gray circles, and the two drug binding sites, Sudlow I & II, are shown by an arrow. The four bottom-right panels show what percentage of the AEBSF sites in each cluster **(C)** is in each HSA domain, **(D)** is on each amino acid residue, **(E)** has each secondary structure, and **(F)** the average SASA score of each cluster. Only SASA scores between C1 and C2 are statistically different (p = 0.019).

When each AEBSF modification intensity is compared between RA and non-RA subjects in the C1, C2, and C3 clusters, all appear to be less accessible in RA HSA. Statistical analysis reveals that three C1 sites (Y263, K359, and H367, in subdomain IIB) and two C2 sites (Y401 and Y497, in subdomain IIIB) have p-values below 0.05, indicating potential RA-specific binding sites **(Table 1, indicated in Figure 6A)**. These significant sites are labeled in **Figure 6A**. As explained for **Figure 5C**, we do not expect potential binding partners at these C1 and C2 sites to increase HSA thermal stability. This hypothesis is strengthened by the fact that most C1 and C2 sites (particularly the significant ones) appear on the more outer surfaces of HSA **(Figure 6A)** and potentially mobile helices **(Figure 6E)**, making them less likely to have a significant impact on overall HSA stability. On the other hand, C3 sites appear to be more concentrated to inner folds of HSA, where a large number of core interactions would need to be broken during denaturation. Interestingly, no individual C3 sites show statistically significant differences between RA and non-RA groups **(Table 1)**, but as a block there is an enrichment in non-RA subjects with high C3 sites **(Figure 5C)**. This suggests that the HSA structure is modified by dynamic surface interactors like other proteins, rather than covalently cross-linked molecules.

**Table 1:**
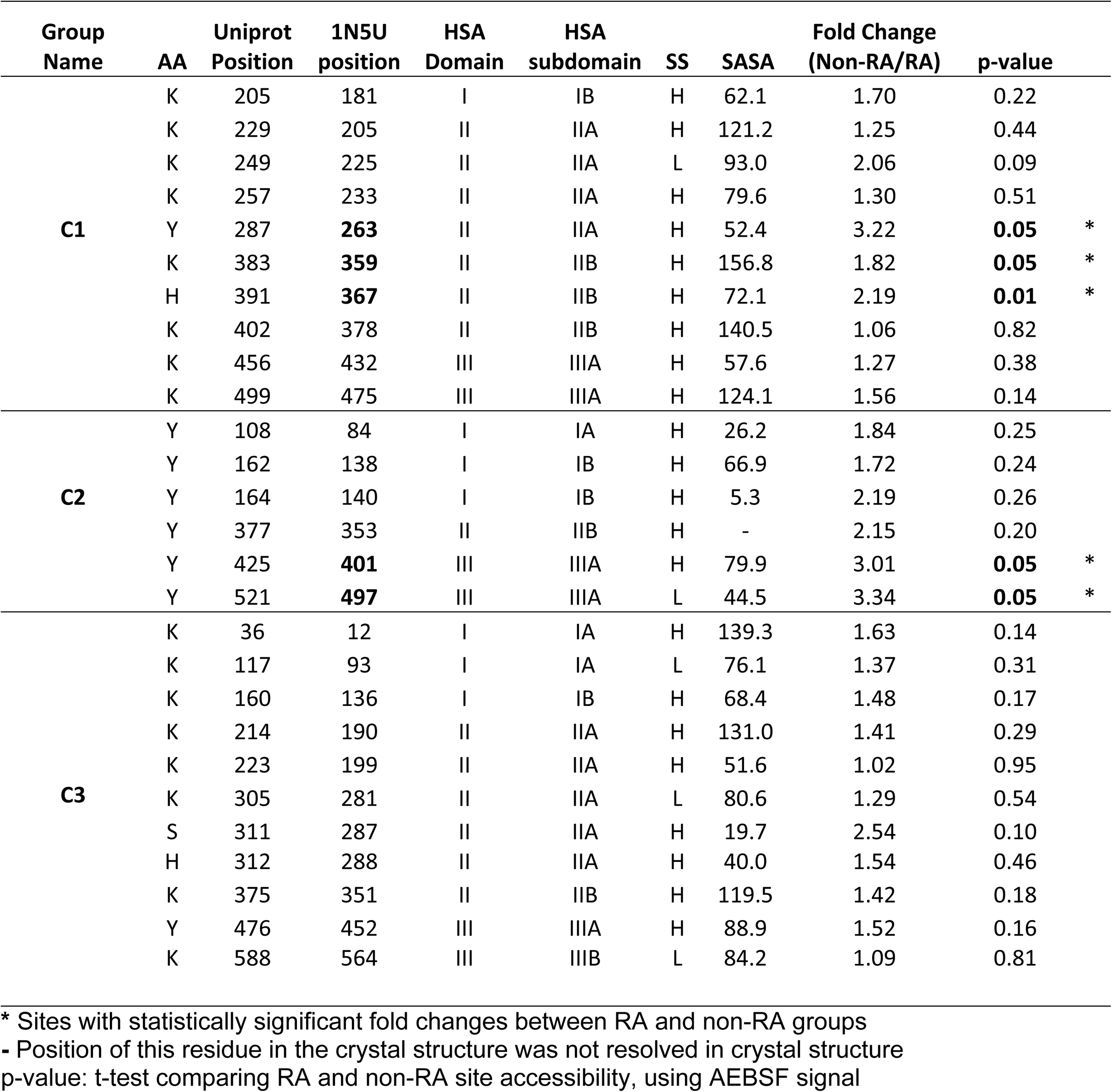
Each of the AAmod sites for groups C1, C2, and C3.

| Group Name | AA | Uniprot Position | 1N5U position | HSA Domain | HSA subdomain | SS | SASA | Fold Change (Non-RA/RA) | p-value |  |
| --- | --- | --- | --- | --- | --- | --- | --- | --- | --- | --- |
| **C1** | K | 205 | 181 | I | IB | H | 62.1 | 1.70 | 0.22 |  |
|  | K | 229 | 205 | II | IIA | H | 121.2 | 1.25 | 0.44 |  |
|  | K | 249 | 225 | II | IIA | L | 93.0 | 2.06 | 0.09 |  |
|  | K | 257 | 233 | II | IIA | H | 79.6 | 1.30 | 0.51 |  |
|  | Y | 287 | **263** | II | IIA | H | 52.4 | 3.22 | **0.05** | * |
|  | K | 383 | **359** | II | IIB | H | 156.8 | 1.82 | **0.05** | * |
|  | H | 391 | **367** | II | IIB | H | 72.1 | 2.19 | **0.01** | * |
|  | K | 402 | 378 | II | IIB | H | 140.5 | 1.06 | 0.82 |  |
|  | K | 456 | 432 | III | IIIA | H | 57.6 | 1.27 | 0.38 |  |
|  | K | 499 | 475 | III | IIIA | H | 124.1 | 1.56 | 0.14 |  |
| **C2** | Y | 108 | 84 | I | IA | H | 26.2 | 1.84 | 0.25 |  |
|  | Y | 162 | 138 | I | IB | H | 66.9 | 1.72 | 0.24 |  |
|  | Y | 164 | 140 | I | IB | H | 5.3 | 2.19 | 0.26 |  |
|  | Y | 377 | 353 | II | IIB | H | - | 2.15 | 0.20 |  |
|  | Y | 425 | **401** | III | IIIA | H | 79.9 | 3.01 | **0.05** | * |
|  | Y | 521 | **497** | III | IIIA | L | 44.5 | 3.34 | **0.05** | * |
| **C3** | K | 36 | 12 | I | IA | H | 139.3 | 1.63 | 0.14 |  |
|  | K | 117 | 93 | I | IA | L | 76.1 | 1.37 | 0.31 |  |
|  | K | 160 | 136 | I | IB | H | 68.4 | 1.48 | 0.17 |  |
|  | K | 214 | 190 | II | IIA | H | 131.0 | 1.41 | 0.29 |  |
|  | K | 223 | 199 | II | IIA | H | 51.6 | 1.02 | 0.95 |  |
|  | K | 305 | 281 | II | IIA | L | 80.6 | 1.29 | 0.54 |  |
|  | S | 311 | 287 | II | IIA | H | 19.7 | 2.54 | 0.10 |  |
|  | H | 312 | 288 | II | IIA | H | 40.0 | 1.54 | 0.46 |  |
|  | K | 375 | 351 | II | IIB | H | 119.5 | 1.42 | 0.18 |  |
|  | Y | 476 | 452 | III | IIIA | H | 88.9 | 1.52 | 0.16 |  |
|  | K | 588 | 564 | III | IIIB | L | 84.2 | 1.09 | 0.81 |  |

\* Sites with statistically significant fold changes between RA and non-RA groups - Position of this residue in the crystal structure was not resolved in crystal structure p-value: t-test comparing RA and non-RA site accessibility, using AEBSF signal

The most significantly altered C3 amino acid residue (in terms of surface accessibility) is S287. Compared to RA subjects, non-RA subjects had 2.54 times has much AEBSF modification at the S287 site (p = 0.10). In the PDB structure, S287 already appears quite buried in HSA **(Figure 6)**, and its SASA score is only 19.7, the lowest of all C3 sites **(Table 1)**. The next two most altered C3 sites are K12 and Y452 (near S287) – sites at which non-RA subjects have 1.63 and 1.52 times as much AEBSF modification (p-values are 0.14 and 0.16, respectively). Seven of the C3 sites (including these top 3) are in a small, localized area (oval shaped magnification in **Figure 6A**) in domain I that could be a plausible binding interface with RA-specific interactors. Binding interactions could increase thermal stability of HSA and reduce the surface reactivity of these sites.

## CONCLUSIONS

In agreement with literature on other diseases (30, 32), we found that the HDC of serum are characteristically shifted (shown by a decreased first/second peak ratio) in all subjects experiencing inflammatory symptoms. Interestingly, RA subjects displayed an even lower peak ratio compared to non-RA subjects, suggesting a more pronounced HDC shift. In comparison, all 15 of the healthy control subjects used by Garbett et al. (22) fall into the HPR group (peak ratio > 1.00, Table 2), 54.5% of non-RA subjects, but only 18.5% RA subjects fell into the HPR group. Our data is consistent with the literature showing that concentrations of the top 8 proteins do not change significantly during RA or other cases of inflammation (61). Our data supports the proposed mechanism in **Figure 2B**, that an increase in HSA stability (5–15°C increase in melting temperature for ∼10% of HSA) would be a more plausible explanation for difference between non-RA and RA HDCs. CRP, a biomarker known to defend against infectious agents and play a significant role in the inflammatory response (4, 62), is the only protein with a significantly different concentration among both comparisons. Both groups are expected to have elevated CRP concentrations, but relative concentrations are less elevated in both RA and LPR groups, compared to non-RA and HPR groups. At the same time, we observed that surface reactivity in the C1, C2, and C3 clusters are less accessible in HSA of RA subjects compared to non-RA subjects, but only sites in the C3 cluster appear to also be less accessible among the LPR group.

**Table 2:** A summary of non-RA and RA subjects versus literature for healthy subjects.

| Quantitative metric | Healthy (literature) | Non-RA (experimental) | RA (experimental) |
| --- | --- | --- | --- |
| HDC ratio | Baseline: $1.59 \pm 0.04$ (22) | $1.00 \pm 0.23$ | $0.83 \pm 0.16$ |
| Peak Ratio Group | 100% HPR (22) | 54.5% HPR<br>45.5% LPR | 18.5% HPR<br>81.5% LPR |
| Concentration of top 8 abundant proteins | Baseline, similar to RA and symptomatic patients (61) | No change compared to RA | No change compared to non-RA |
| CRP concentration | Baseline, low compared to non-RA and RA (4) | High | Elevated, but lower than non-RA |
| C1 | <u>Binding previously observed:</u><br>K205 (79)<br>K225 (80)<br>K233 (81)<br>K359 (82)<br>H367 (81)<br>K378 (81)<br>K432 (80, 86, 88) | More surface reactivity compared to RA; significant on sites Y263, K359, H367 | Lower surface reactivity compared to non-RA; appears to be associated with a higher thermal stability |
|  | <u>Reactivity previously observed:</u><br>K181 (91), K263 (92), K475 (91) |  |  |
| C2 | <u>Binding previously observed:</u><br>Y84 (83)<br>Y138 (84, 86)<br>Y353 (85)<br>Y401 (86)<br>Y497 (87) | More surface reactivity compared to RA; significance on sites Y401, Y497 | Lower surface reactivity compared to non-RA; appears to be associated with a higher thermal stability |
|  | <u>Reactivity previously observed:</u><br>Y140 (93) |  |  |
| C3 | <u>Binding previously observed:</u><br>K12 (80)<br>K136 (89)<br>K190 (77, 80, 86)<br>K199 (77, 80, 86, 88)<br>K281 (78)<br>S287 (77)<br>H288 (90)<br>K351 (79, 80, 82, 86)<br>Y452 (88) | More surface reactivity, suggesting a lost binding interface that exposes K12, K199, K281, S287, H288, and Y482 | Lower surface reactivity compared to non-RA; appears to be associated with a higher thermal stability |
|  | <u>Reactivity previously observed:</u><br>K93 (91), K564 (91) |  |  |

These findings suggest a model consistent with **Figure 2B**. CRP, produced predominantly by hepatocytes in response to stimulation by IL-6, is known to be a promiscuous interactor and recruiter of proteins (63, 64). For example, CRP binding to immunoglobulin Fc gamma receptors (FcgR) promotes the production of proinflammatory cytokines leading to an amplification loop of inflammation. (62). It is possible that during inflammation in non-RA subjects, high levels of CRP associate with HSA binding proteins, sequestering them from HSA. Since CRP is decreased in RA subjects, these potential HSA interactors would be more available to bind HSA, specifically at the C3 site surface interface (in domain I of HSA) shown in Figure 6A, increasing HSA stability, and reducing surface accessibility.

The BioGRID 4.4 interactome database shows 349 known interactors for HSA and 72 interactors for CRP. Three of these proteins are common interactors to both HSA and CRP: 1-acylglycerol-3-phosphate O-acyltransferase 1 (AGPAT1), complement factor H (CFH), and fibronectin 1 (FN1). Some studies have shown brief details about CRP’s relationship with AGPAT1 (65, 66), but extensive studies have shown the specificity and details of CRP’s interactions with CFH (67–70) and FN1 (71–76). The specific location of these proteins’ interactions with HSA is less understood. Nevertheless, most of the C3 modification sites such as those in the plausible domain I binding pocket (K12, K199, K281, S287, H288, and Y482) are known to be high affinity binding sites for drugs or other protein interactors (77–90). **Table 2** shows which of the C1, C2, and C3 sites have been associated with ligands, drug, or protein binding sites in previous studies (77–90). For all other sites, reactivity has still been observed (91, 92, 93).

In an effort to better understand RA pathology and potential HSA interactors, future RA studies should look for potential binding partners by extracting lipids and other cargo from purified serum HSA. Other directions to explore the mechanisms that increase HSA stability are: (1) Using more specific surface modifications or chemical crosslinking reagents to carry out more in-depth surface probing of HSA, allowing more specific information about HSA binding partners and conformation changes, and (2) comparing HSA and CRP protein binding partners in RA and non-RA patients using immunoaffinity purification together with mass spectrometry to understand how a relative decrease in CRP could be contributing to HSA protein interactors. Future research into HSA and other related proteins will continue to enhance our understand RA-specific pathology and give insights into the development of, and potential treatments for, RA.

## METHODS

### Heat Denaturation Curves

50 blood serum samples were taken by ARUP Laboratories to the TA Instruments application lab. Samples were prepared in random order for Nano Differential Scanning Calorimetry DSC measurements by first filtering with a 0.45-micron filter. After being degassed, 40 µL of the blood serum was diluted with 960 µL of buffer. The buffer used for dilution was 10 mM phosphate-buffered saline (PBS) (138 mM NaCl, 2.7 mM KCl at pH 7.50). Samples were refrigerated at 4 °C until Nano DSC scans were made. Samples were prepared ten at a time and loaded into the Nano DSC autosampler at 5 °C. Samples were scanned from 20 ° to 110 °C at 1 ⁰C/min after a 600 second equilibration period after loading. The remainder of the undiluted serum samples were taken to BYU for MS analysis to look for changes in protein concentration, as well as PTM frequency and location.

### Calculating the Peak Values

Calorimetry experimental results were first corrected for the instrument baseline by subtracting a buffer injection control. Nonzero baselines were then corrected by applying a linear baseline fit. Scans were finally normalized for the concentration of protein injected. We then looked at the raw HDC curve between 25 and 100°C, setting the minimum of each HDC as 0 and the maximum as 100. This allowed us to take the peak ratio from two positive values. The first peak value (HSA peak) was measured at 63°C, and the second peak value (Ig peak) was measured at 71°C. The HSA/Ig peak ratio was then calculated.

### Protein Digestion

The serum samples were denatured with 6M guanidine chloride (GdmCl) in 100mM Tris/HCl (pH 8.5) and protease inhibitor (Sigma-Aldrich, cat #: P8340), then spun at 21,000xg for 20 minutes at 4 °C to remove insoluble cell contents. The supernatant, which contains soluble proteins, was then transferred into new tubes. The BCA assay (Thermo Fisher Scientific cat #: 23227) protocol was followed to measure the protein concentration in each sample. 1.5 µL serum, which contained about 50 µg of protein, was diluted to 50 µL in 1X PBS, and combined with 100 µL 6 M GdmCl. Each sample was transferred into a new tube, then 1.2 µL of 200 mM dithiothreitol (DTT, >99% sigma # D-5545) in water was added (final concentration 5mM) and the mixture was incubated at 55°C in a sand bath for 15 minutes. The mixture was then cooled for 5 minutes to reduce disulfide bonding. We then added 3.8 µL of 200 mM freshly made iodoacetamide (IAM, 97% sigma # I-670-9) in water (final concentration 15 mM) and incubated for 1 hour at room temperature in the dark to alkylate the reduced proteins.

Next, samples were put onto 30 kDa centrifugal filters and spun at 14,000 g for 10 minutes. Then 100 µL 6M GdmCl in 100mM Tris/HCl (pH 8.5) was added, and the samples were spun at 14,000xg. This was repeated twice. Then 100 µL 25 mM ammonium bicarbonate (ABC) was added, and the samples were spun again at 14,000 g, this was repeated twice. Next, we emptied and cleaned the collection tube with ddH_2_O three times and 100 µL 25 mM ABC was added to the top of the filter.

MS trypsin (Promega gold MS sequencing grade Trypsin #V5111) was added to the solution above the filter in a 1:50 (w/w) trypsin/protein ratio and the samples were incubated at 37°C overnight on a shaker. After that, each sample was quenched with 300 mM phenylmethylsulfonyl fluoride (PMSF, final concentration 1 mM). Samples were then centrifuged at 14,000xg for 30 minutes, 100 µL of 25 mM ABC was added, and the samples centrifuged again at 14,000 g for 30 minutes. The filtrate was collected in mass spec vials, dried with a Speedvac, and resuspended in 3% ACN, 0.1% FA to 1 µg/µL.

### Mass Spectrometry Acquisition for Proteomics

Data for the 50 samples was acquired in a randomized order. Digested peptides were separated on a Polaris-HR-C18 HPLC chip in a chip cube nano spray source using an Agilent 1260 HPLC followed by positive ESI and mass detection using an Agilent QTOF mass spectrometer (6530B). The mobile phases consisted of MS grade 3% acetonitrile, 0.1% formic acid for Buffer A; and 97% acetonitrile, 0.1% formic acid for Buffer B. A 50-minute gradient was run at 0.3µL/min flow rate: 0%-5% B Buffer (0-0.5 minutes), 5%-30% B Buffer (0.5-27 minutes), 30%-95% B buffer (27-28 minutes), 95% B Buffer (28-31 minutes), 95% - 5% B buffer (31 - 33 minutes), 5% - 95% B buffer (33 - 35 minutes), 95% - 0% B buffer (35 – 46 min), 0% B (36-49 min). Auto MS/MS fragmentation using variable collision energy determined by ion mass from 290-1700 m/z at 4 spectra/s rate and 250ms/spectrum time, and with an isolation width around 4 m/z. The auto MS/MS method selected precursor ions that were above 2500 counts and have charge state 2 and above for fragmentation. MS/MS scan range 100-1700 m/z, and 10 max processors allowed per cycle. The same spectra were excluded from the MS/MS selection for 0.2 min. This prevented continual acquisition of the same m/z and allowed for other, less abundant species to be acquired by the mass spectrometer.

### Protein Identification and Quantification

Protein identification and quantification were performed with two programs. The first was Protein Prospector developed in the University of California San Francisco Mass Spectrometry Facility, funded by NIH National Institute for General Medical Sciences. The second one was PEAKs Studio 8.5, developed Bioinformatics Solutions Inc. Both programs compared peptide fragmentation against the SwissProt human database downloaded in August 2017 with the following parameters: monoisotopic for precursor mass search type; semispecific for digest mode, 3 missed cleavage allowed; 20 ppm for parent mass error tolerance; 0.5 Da for fragment mass error tolerance; 3 max variable PTM per peptide allowed, with carbamidomethlyation as fixed modification, and oxidation, Pyro-glu from Q and other 9 customized PTM as variable modification (detailed listed in **Supplemental Data 4**) in PEAKS DataBase step; 311 built-in ptm was used in the PEAKS PTM step; 20ppm mass error tolerance and 3 min retention time shift tolerance were used in the label free quantification step. The raw data are available for download at the chorusporject.org (project ID: 1739, experiment ID:3632).

### Protein structure analysis

Some analyses performed with UCSF Chimera, developed by the Resource for Biocomputing, Visualization, and Informatics at the University of California, San Francisco, with support from NIH P41-GM103311.

### Inferno Hierarchical Clustering and Heatmap analysis

The heatmap for AEBSF on HSA was created using InfernoRDN created by Pacific Northwest National Laboratory (PNNL, 53). PTM sites were identified and quantified using PEAKS Studio. PTM sites that were at least present in 12 samples were included in Heatmap generation. Files were then loaded into InfernoRDN and Log2 transformed to reduce the noise of outliers in later analysis. A dual-clustered Heatmap was generated with the standard Euclidean modeling parameters. The hierarchical order output was then used to determine the most changed PTM sites between samples and subsequent PTM site groupings. Additionally, the dual-clustering setting allowed for groups to be observed across samples which were statistically examined for correlation with RA diagnosis.

## Abbreviations

DSC: Differential scanning calorimetry
HDC: Heat Denaturation Curves
MS: mass spectrometry
HSA: human serum albumin
RA: rheumatoid arthritis
PTM: post translational modification
RF: rheumatoid factor
ACPA: anti-citrullinated peptide antibodies
AKA: anti-keratin antibody
HDL: high-density lipoprotein
CCP: cyclic citrullinated peptide
ARUP: Associated Regional and University Pathologists
HAPT: Haptoglobin
Ig: Immunoglobulin
LPR: low peak ratio
HPR: high peak ratio
LC–MS/MS: Liquid chromatography–tandem mass spectrometry
QTOF: Quadrupole Time-of-Flight
CRP: C-reactive protein
VDBP: Vitamin D binding protein
AAmod: amino acid modification
AEBSF: aminoethylbenzenesulfonylflouride
SASA: surface accessible surface area
GdmCl: guanidine chloride
DTT: dithiothreitol
IAM: Iodoacetamide
PBS: phosphate-buffered saline
ABC: ammonium bicarbonate
PMSF: phenylmethylsulfonyl fluoride
PNNL: Pacific Northwest National Laboratory

## ACKNOWLEDGMENT

We gratefully acknowledge ARUP laboratories (Salt Lake City, UT) for providing anonymized serum samples after the RA panel was finalized; TA Instruments Application lab (Lindon, UT) for access to the DSC instrumentation and expertise. BYU undergraduate research awards supported JCH, CT, DHP, MH, SA, NRZ. Research reported in this publication was supported by the Fritz B. Burns Foundation and the National Institute on Aging of the National Institutes of Health under Award Number R01AG066874 to JCP. The content is solely the responsibility of the authors and does not necessarily represent the official views of the National Institutes of Health.

## SUPPORTING INFORMATION

### Supplemental Figures

- Supplemental Figure 1-All HDCs per group
- Supplemental Figure 2-Average HDCs with Deviation
- Supplemental Figure 3-HDC simulation
- Supplemental Table 1-Proteins have Significant Changes
- MS2 spectra for PTM on HSA

### Supplemental Data 1 (.xlsx): Samples Information and Classification

- *<u>Sample Information:</u>* Includes sample name, age, gender, CCP/RF values, and clinical RA diagnosis. This was provided by ARUP laboratory.
- *Classification in each experiment:*

- HDC Peak Ratio: The ratio of the HSA peak at 61°C to the Ig peak at 73°C
- HDC group: the group is assigned based on HDC peak ratio. The HPR has peak ratio > 1.00, the LPR has peak ratio < 1.00
- PTM group: the group is assigned based on hierarchical order of the Inferno Heatmap in **Figure 4B**. It is used for analysis shown in **Figure 4C**.
- *<u>Experiment raw data filename:</u>* the directory of the HDC and MS filenames in relation to the sample name. The MS raw files are available for download at the chorusporject.org (project ID: 1739, experiment ID:3632). The HDC raw files are available upon request.

### Supplemental Data 2 (.zip): HDC Results/Simulation

- The zip file includes 47 HDC results (.csv) exported from DSC raw files. Each HDC result file contains seven measurements for each DSC run: temperature (°C), power (µW), time(s), pressure (atm), scan rate (°C/min), analysis data (excess molar heat capacity (Cpex)), and corrected data (normalized Cpex, normalization explained in the Methods section).

### Supplemental Data 3 (.xlsx): Protein Quantification

- **protein-peptide:** the peptide area exported from LFQ from PEAKs Studio for the 49 samples. This is used for protein quantification, and PTM analysis.
- **Filter:** The filters applied for LFQ analysis.

### Supplemental Data 4 (.xlsx): PTM Results for HSA

- **PTM Results for HSA**

- **ProteinProspector:** Lists the PTM search result on HSA.

- *<u>Peptide information:</u>* Lists peptide sequence, peptide start position, peptide end position, peptide theoretical mass, precursor m/z, precursor mass, and precursor mass.
- *<u>First/Second Modification:</u>* Lists the amino acid, the position, the mass shift value of the modification, as well as SLIP score (a quality merit of the modification)
- *<u>Hit:</u>* List if the modified peptide is observed in a sample. If it is present, the sample name is record in the same row of the peptides. The analysis only returns present or not, thus PEAKs studio is used for further quantification.
- **Customized PTM search**: Lists the name, m/z shift, modified AA for the PTM put in the database search step of PEAKs studio analysis.
- **Albu_ptm profile:** The data from PTM profile of PEAKs Studio SPIDER analysis. It lists all PTM observed on HSA, and the modification site. The peptide sequence window, the modified amino acid (AA), the occurrence of the same modified AA in the dataset, the modified site on the protein, the occurrence of the same modified site in the dataset, best-10logP, best ion intensity (%), and number of hit across 49 samples.

### Supplemental Data 5 (.xlsx): AEBSF sites on HSA

- **T-test:** Lists t-test results between RA and non-RA subjects, as well HPR and LPR subjects, showing significance between RA and non-RA subjects in each comparison. The average intensity of the sum of all of AEBSF modification sites on HSA of each sample, for RA and non-RA, as well as HPR and LPR, is shown. A t-test was also performed between groups for modification sites in clusters 1, 2, and 3.
- **AEBSF_HSA (site):** This dataset lists the structure characteristic of the 41 AEBSF sites on HSA, including association with the cluster groups from Figure 4B, HC order from Inferno, site position from Uniprot, site position from PDB ID 1N5U, HSA domain, HSA subdomain, amino acid (AA), secondary structure (SS), surface accessible surface area (SASA) score, number of peptides used for the quantification of the site, peptide sequence, and the intensity from each sample. (The intensity here used the area from supplemental data 3. Only peptides from HSA and with AEBSF modifications are retained. The area of peptides that have same AEBSF modification site are combined (the number of peptides is used for combination is listed in column #peptide combined). After consolidation, site that have less than 12 hits are removed. Note that the sequence/start/end for sites that used more than 1 peptides are just representative. The intensity is also used for Inferno analysis).

**AEBSF_HSA_(cluster site) :** A list of all samples, with their groups (RA/non-RA and HPR/LPR), as well as the intensity sum of AEBSF modification for each modification site in cluster C1, C2, C3 on HSA.

